# Translation Efficiency Impacts Phage Lysis Timing and Its Precision in Single Cells

**DOI:** 10.1101/2025.05.14.654169

**Authors:** Sherin Kannoly, Kevin Singh, Naseerah Juman, Zubaida Marufee Islam, Iñigo Caballero Quiroga, Abhyudai Singh, John J. Dennehy

## Abstract

The timing of host cell lysis is a fundamental life history parameter in bacteriophages as it presents an evolutionary trade-off between maximizing intracellular phage replication and optimizing transmission to new host cells, ultimately determining viral fitness in dynamic bacterial populations. In the bacteriophage λ, lysis time is dependent on the expression of a timekeeper protein, holin, which accumulates in the *Escherichia coli* inner membrane. Cell lysis is triggered when the membrane concentration of holin crosses a critical threshold level. In the present study, we investigated the effects of the rate of holin translation on lysis timing and its precision. We show that modulating holin translation efficiency through genetic modifications and antibiotic treatment alters bacteriophage lysis timing and its precision, providing insight into how phages optimize the evolutionary trade-off between replication and transmission. Reducing ribosomal binding affinity decreased holin expression and delayed lysis, while optimizing the Shine-Dalgarno sequence enhanced translation and accelerated lysis. Surprisingly, very low tetracycline concentrations may have improved translation efficiency and hastened lysis, whereas higher doses predictably delayed it. In all cases, longer lysis times corresponded with decreased timing variability. A model that incorporates stochastic gene expression with additional stochasticity in the initiation of holin expression was sufficient to explain the data. Thus, we demonstrated that modifying the rate of holin translation can be used as an evolutionary strategy to achieve optimum lysis timing.

**Importance:** Our prior work demonstrated that phage lysis timing is calibrated to optimize fitness based on environmental conditions affecting host availability and phage replication. We identified an optimal threshold—the critical holin concentration—that minimizes lysis time variability. Genetic manipulation of the holin gene altered the lysis threshold and generated phages with non-optimal early and late lysis times. The current study reveals that adjusting holin translation rates offers another mechanism for modifying lysis timing. Phages therefore have multiple means to adjust the timing of host lysis in order to maximize fitness under different selective pressures.

## Introduction

We used the bacteriophage lambda (λ) to study the timing of host cell lysis and its precision. At the end of the infectious life cycle, phage λ induces the rupture of the host cell (lysis) to release its own progeny. Lysis is triggered when the λ-encoded regulatory protein “holin” reaches a critical threshold concentration in the inner membrane of its host *Escherichia coli*. Upon reaching this threshold, holin nucleates to form large lesions or pores in the inner membrane triggering the instantaneous rupture of the cell (1–3). Thus, holin acts as a molecular timekeeper, precisely regulating the timing of cell lysis to ensure optimal release of phage progeny (1,4). Lysis time directly impacts phage progeny production (i.e., the burst size) (5). The longer the lysis time, the larger the burst size (6). Thus, lysis time directly influences phage fecundity. Previously, we showed that there exists an optimum range of lysis times that maximizes phage production under quasi-steady state conditions, where cells and phages are regularly washed out (7).

Using simple microscopy techniques, λ-induced cell lysis can be readily visualized and recorded at the single-cell level, allowing for accurate estimation of both the mean lysis time and its associated noise (8). In a previous study, we used site-directed mutagenesis of the holin gene to generate λ mutants that varied in their lysis times ranging from 30 to 190 min (9). The wild type λ showed a remarkable precision in lysis timing with a coefficient of variation of less than 10%. However, the mutant strains exhibited considerable variability in lysis timing due to differences in the critical holin concentration needed to initiate lysis. In our model, “threshold concentration” represents holin’s capacity to trigger cell lysis. Amino acid substitutions in holin modify this threshold either by changing the concentration required to initiate lysis or by altering the efficiency of membrane incorporation.

The aim of this study was to investigate the effect of manipulating the rate of holin translation on lysis timing and its precision, while leaving the holin threshold (i.e., protein amino acid sequence) unchanged. Based on previous studies, we identified nucleotides in the translation regulatory regions that impacted holin translation efficiency (10). These regions play important roles in ribosome binding and the formation of translation initiation complexes. Nucleotide substitutions that strengthen or weaken ribosome binding will alter the rate of holin translation.

In prokaryotes, the translation start site begins with a single AUG start codon encoding the amino acid methionine (Met) (11). Usually, this start codon is positioned 6−10 nucleotides downstream of a Shine-Dalgarno (SD) sequence. This position allows for optimal binding of 30S ribosomal subunit to properly initiate translation by distinguishing the start codon from the AUG codons internal to the reading frame of a polycistronic mRNA. The λ *S* gene encodes two protein products, holin and antiholin in the same reading frame (12–14). Antiholin is able to dimerize with holin and prevent it from participating in hole formation, which delays the lysis time, but the biological significance of this inhibition remains unclear (15,16). The 5′ end of the gene contains two AUG start codons separated by a single codon encoding lysine. Chang et al. investigated the effects of mutations in the holin translation initiation region on lysis timing (10). Mutational studies of the translational initiation region found that four elements regulate translation initiation from the two holin gene AUG start codons (Met1 and Met3): 1) an upstream near-consensus SD sequence (GGGGG from nucleotide (nt) −12 to −7) serving as the 16S RNA-binding site for expression from Met1 (which results in the production of antiholin), 2) an unusual SD sequence (UAAG from nt −6 to −3) serving the Met3 start codon (which results in the expression of holin), 3) an upstream stem-loop structure (nt −21 to −7) controlling ribosomal access to the GGGGG sequence, and 4) a downstream intragenic stem-loop (nt 31 to 47) structure (Fig. 1A) (14). The two stem-loop structures appear to complement each other in regulating the partition of translational initiation between the two start codons in a 1:2 ratio favoring holin (14).

**Figure 1.**
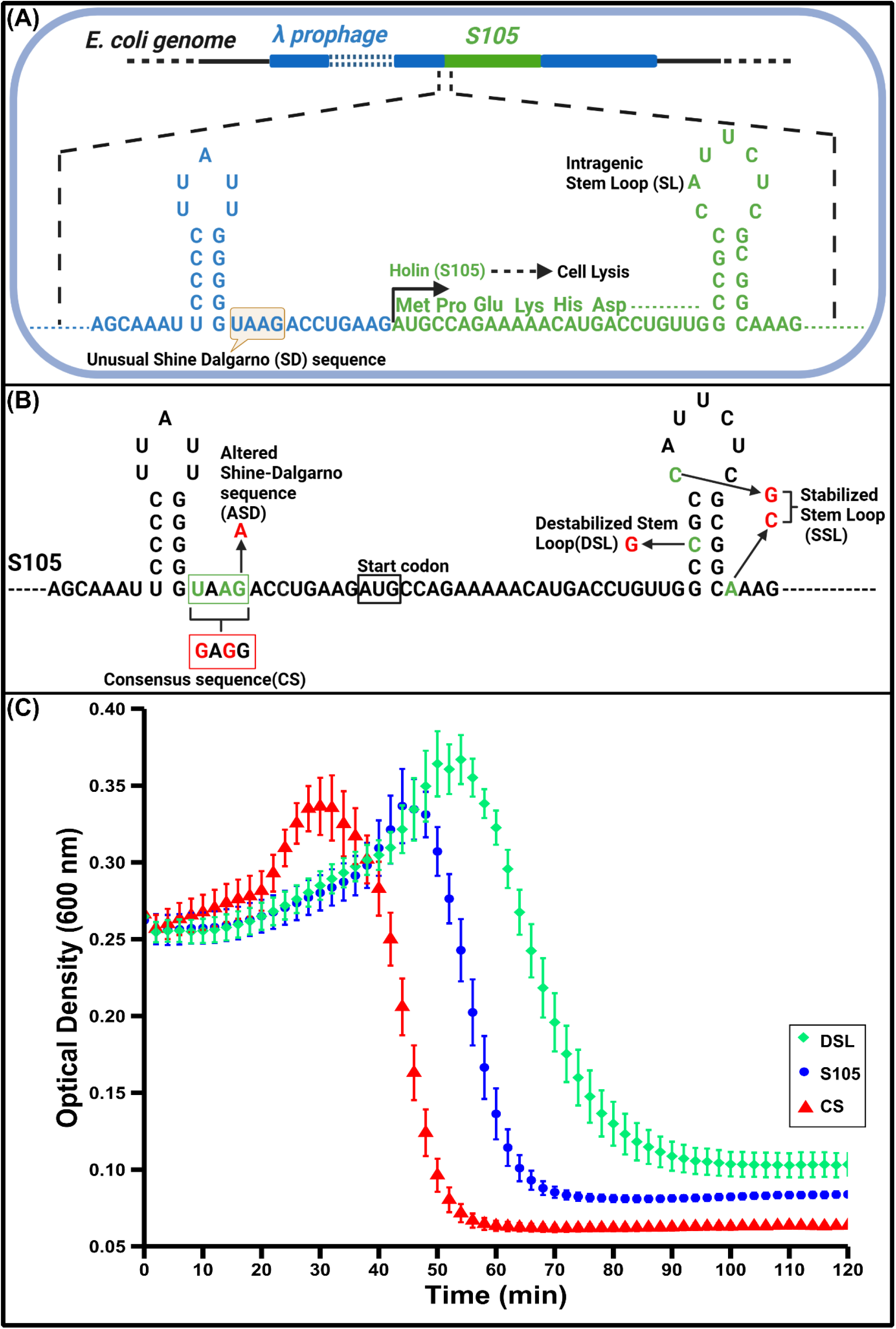
Structure of the holin translation region in the S105 lysogenic strain. (A) The sequence of the 5′ region of mRNA is shown with stem-loop RNA secondary structures, which were previously characterized in vivo and in vitro (10,13,14). (B) The N-terminal amino acid sequence of holin (S105) starting from methionine (Met) is shown above the mRNA. The intragenic stem loop (SL) has been shown to regulate the formation of translation initiation complexes at Met (14). The nucleotides targeted for substitutions (in green) in the S105 background are shown. (C) The arrows from these nucleotides indicate sequence changes introduced in the lysogens CS, ASD, DSL, and SSL (in red). The lysogens were grown in LB broth and heat induced to start the lytic cycle. Following induction, the optical densities of the cultures were measured to estimate the lysis time for each lysogen (C). Error bars denote 95% CI (n = 23).

## Materials and Methods

### Bacterial and phage strains

The bacterial strains and plasmids used in this study are listed in Table 1. Bacterial cultures were grown in lysogeny broth (LB) with aeration (200 rpm) at 30°C.

**Table 1.**
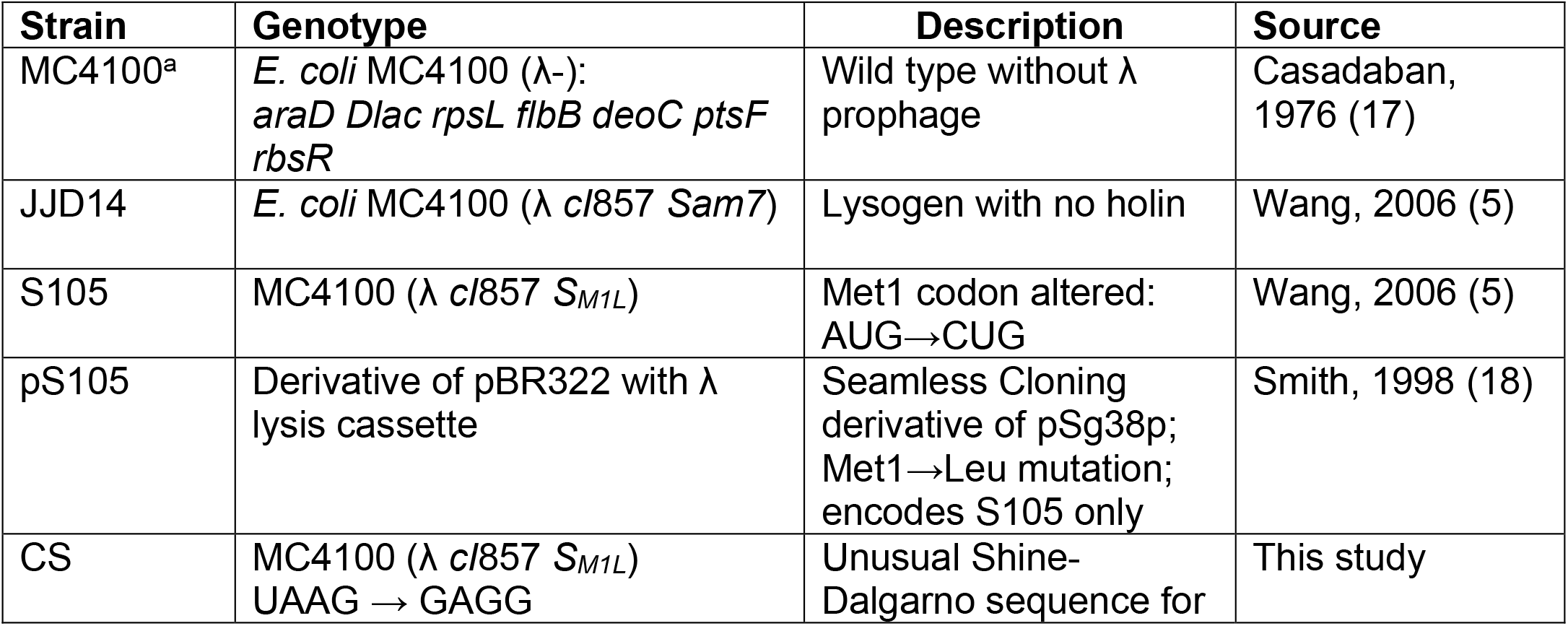
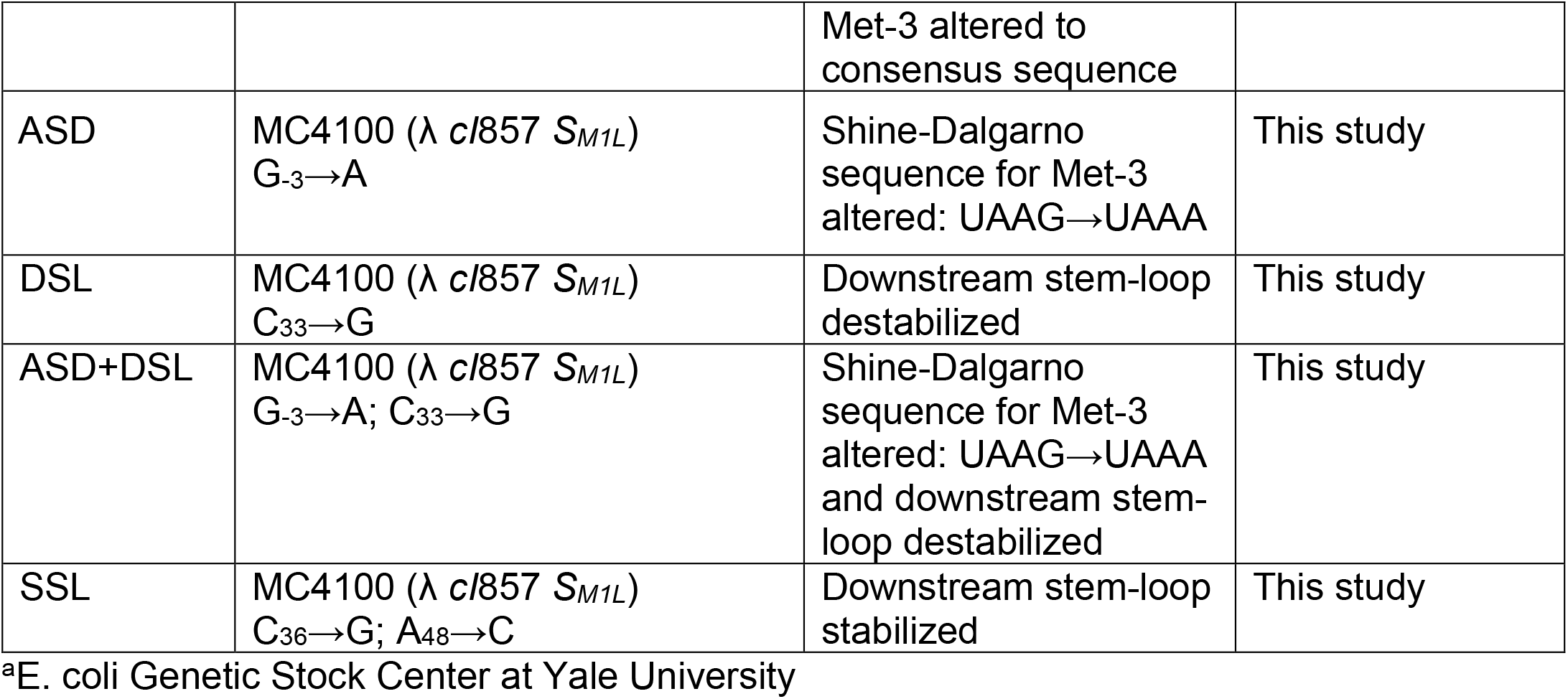
Bacterial strains and plasmids.

### Construction of λ lysogens with mutations that affect holin translation

Site-directed mutagenesis was used to generate λ phages with mutations that affect holin translation (Fig. 1A). Plasmid pS105 carries the λ lysis cassette with the *S105* allele. In the *S105* allele, the Met1 codon of the *S* gene was replaced with a Leu (CUG) codon, thus this phage expresses only the 105 amino acid protein, holin, and not the 107 amino acid protein, antiholin. The plasmid pS105 was used as a template for PCR (Pfu DNA polymerase; Promega, Madison, WI) using megaprimers (Table 2) consisting of 30 to 45-nucleotide homology flanking the altered nucleotides. After PCR, the original pS105 template was digested using DpnI (New England BioLabs, Ipswich, MA) and the mutated plasmids were transformed into *E. coli* lysogen JJD14 (MC4100 [λ cI857 Sam7]) cells. The λ cI857 Sam7 prophage’s holin amino acid sequence is truncated thus its capacity for lysis has been abolished. The transformed cells were spread on LB + Amp (100 µg ml^-1^) plates and incubated at 30°C until colonies were visible. The transformants were grown to an optical density (OD_600_) of 0.2 in 10 mL LB with 100 μg/mL ampicillin at 30°C. The culture was heat shocked at 42°C for 15 min to induce the excision of the prophage from the host chromosome, followed by growth at 37°C until lysis. Plaques that appear on the MC4100 lawn are most likely the results of recombination between the *S* alleles on the plasmid and the prophage and rarely due to reversion of the parental *Sam7* allele. Two plaques were picked, introduced into 10 mL LB with mid-log phase *E. coli* MC4100, and incubated at 37°C for several hours. The resulting cultures were centrifuged at 1,057 RCF for 10 min and the supernatants were passed through a 0.22μm filter to obtain lysates, which were used as templates for PCR amplification using primers that amplified the entire holin gene (S_for_ and S_rev_, Table 2). The resulting PCR product was Sanger sequenced to confirm the presence of the desired nucleotide substitutions. After confirming the correct substitutions, the same lysate was then used to lysogenize *E. coli* MC4100. The resulting lysogens were used for colony PCR with primers S_for_ and S_rev_ and sequenced to confirm the correct mutations were still present.

**Table 2.**
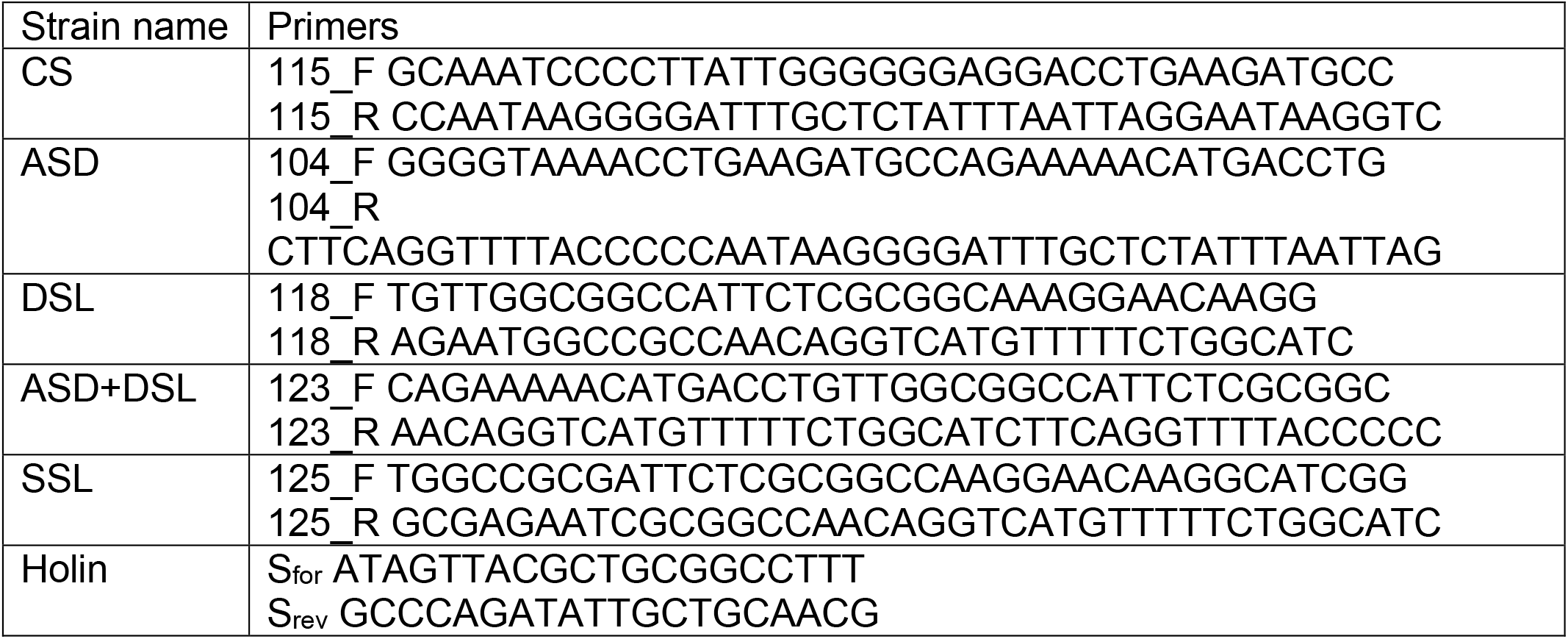
Primers.

### Lysis Time Determination of Batch Cultures Using a Plate Reader

To test the effect of mutations in the translation regulation regions on lysis time, we measured the optical density of cultures following heat induction. Overnight cultures of lysogens were diluted 100-fold in 24-well culture plates (Corning Costar #3738) containing 1 mL LB. The plates were shaken at 30°C for 2.5 h in a plate reader (TECAN Infinite^®^ 200 PRO, Switzerland). For heat induction, the cultures were incubated in a water bath at 42°C for 15 min. Following heat induction, the plates were shaken in a pre-warmed plate reader, which measured the OD_600_ at two-minute intervals.

### Determination of Bacterial Growth Rate

To estimate the bacterial growth rates in the presence of sublethal tetracycline doses, we measured the optical densities (OD_600_) of cultures using a plate reader. Overnight bacterial cultures were diluted 100-fold in 24-well culture plates (Corning Costar #3738) containing 1 mL LB. The plates were shaken at 30°C in a plate reader (TECAN Infinite^®^ 200 PRO, Switzerland) with OD_600_ measurements every ten minutes. The growth rate was estimated using the equation 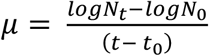, where *μ* is the growth rate and N_0_ and N_t_ are initial and final bacterial numbers, respectively.

### Single-cell Lysis

To estimate single-cell lysis times, we employed the protocol described previously (8). Briefly, lysogens grown overnight at 30°C in LB were diluted 100-fold in LB (with different concentrations of tetracycline wherever appropriate) and grown to OD_600_ = 0.2–0.3 in a 30°C shaking incubator. A 22 mm square glass coverslip was pretreated with 0.01% poly-L-lysine (mol. wt. 150 K–300 K; Millipore Sigma, St. Louis, MO) at room temperature for 10 min. Exponentially growing cells (∼200 µl) were immobilized to the coverslip, which was assembled on a perfusion chamber (RC-21B, Warner Instruments, New Haven, CT). The perfusion chamber was immediately placed on a heated platform (PM2; Warner Instruments, New Haven, CT) mounted on an inverted microscope stage (TS100, Nikon, Melville, NY). Heated LB or LB + Tetracycline at 30°C (Inline heater: SH-27B, dual channel heating controller: TC344B; Warner Instruments, New Haven, CT) was then infused into the chamber. The chamber temperature was increased to 42°C for 20 min, then maintained at 37°C until ∼95% cells were lysed. An eyepiece camera (10X MiniVID™; LW Scientific, Norcross, GA, 10 fps) records video. Lysis times of individual cells were observed and recorded using VLC™ media player. Lysis time was defined as the time required for a cell to disappear after the temperature was spiked to 42 °C (Fig. 2A).

**Figure 2.**
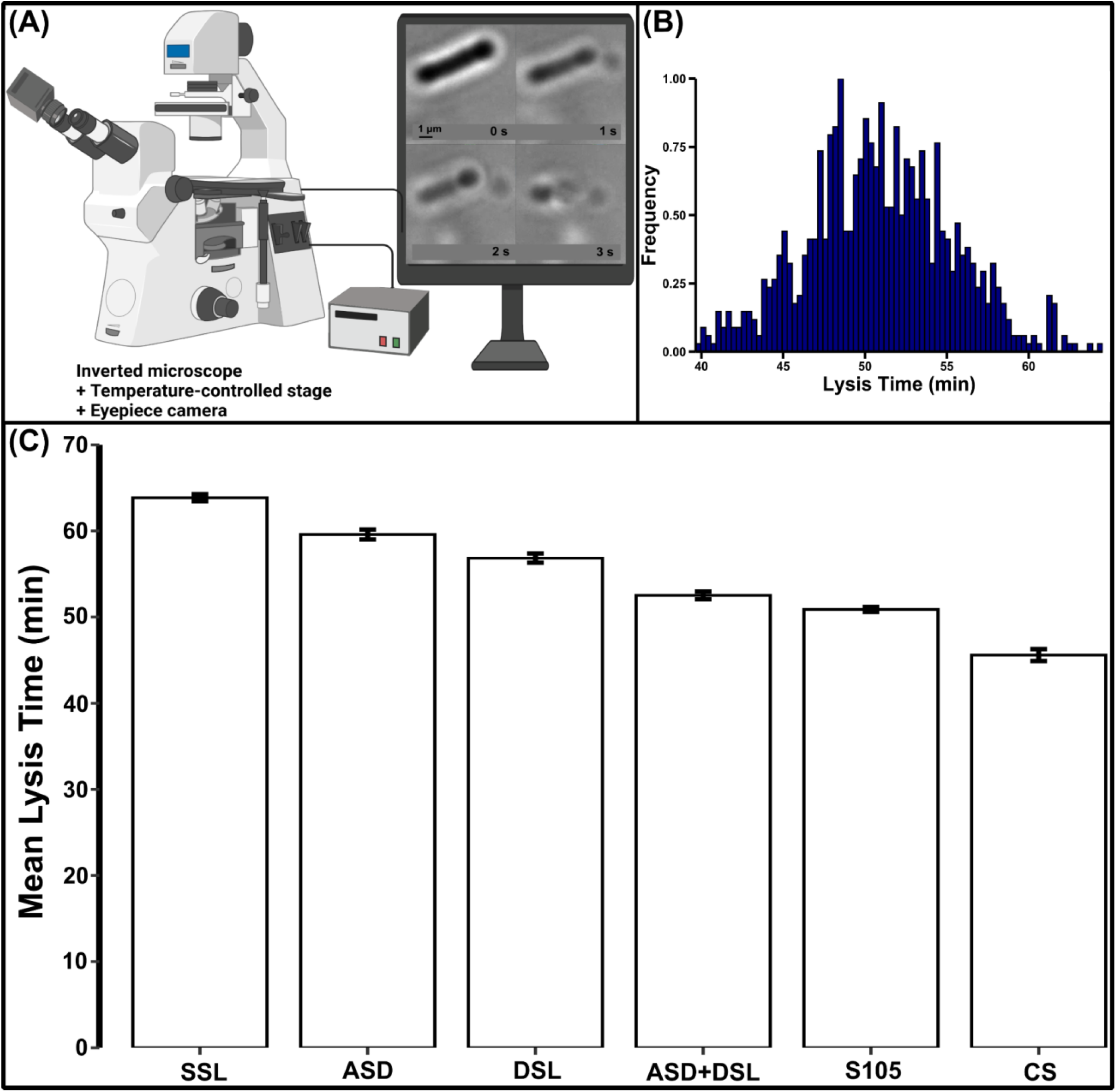
Single-cell experiments to estimate lysis times. (A) Experimental setup for imaging and recording lysis events of single cells. (B) The monitor shows the final three seconds of an induced cell before complete lysis. Single-cell lysis time distributions for S105. (C) Mean lysis times for lysogens with altered translational efficiencies estimated using single-cell lysis times (200–300 cells). Except for the CS lysogen with a shorter lysis time, the ASD+DSL, DSL, ASD, and SSL lysogens show increasingly higher lysis times when compared to that of the S105 lysogen. Error bars represent 95% CIs after bootstrapping (1,000 replicates).

## Results and Discussion

In the present study, we investigated the effects of altered rate of holin translation on lysis timing and its precision. To this end, we employed two different strategies to perturb holin translation. One involved altering regions in the holin mRNA responsible for regulating protein translation. The other involved using the antibiotic tetracycline, which affects translation in bacteria (19–22).

### Effects of Mutations in Holin Translational Regulatory Regions

We used a λ lysogen in which antiholin expression was abolished by introducing the M1L mutation (AUG→CUG) at the Met1 start codon (Fig. 1A). Upon induction, this lysogen expressed only holin (S105), which simplified our model system by removing any antiholin-associated confounding effects (Fig. 1A).

Inspired by Chang et. al, we used site-directed mutagenesis to introduce mutations into the translational regulatory regions of the *S105* gene in a plasmid bearing the lambda lysis cassette (Fig. 1B) (10). The resulting plasmids were used to construct five *E. coli* lysogens with mutations in the translational regulatory region. An S105 lysogen with a near-consensus sequence (CS) serving Met3 start codon was constructed by substituting two nucleotides in the unusual Shine-Dalgarno (SD) sequence (Fig. 1B and Table 1). The unusual SD sequence was also altered by a single nucleotide substitution (ASD, Fig. 1B and Table 1). Strengthening and weakening the downstream stem-loop structure was hypothesized to decrease and increase the formation of initiation complexes at the Met3 start codon, respectively. To test this idea, we attempted to destabilize and stabilize the downstream stem-loop structure by constructing lysogens DSL and SSL, respectively (Fig. 1B and Table 1). Finally, we combined the mutations that alter the SD sequence and destabilize the downstream stem-loop structure (ASD+DSL).

The rate of holin expression directly impacts λ lysis times. A higher rate of holin expression will allow a faster crossing of the lysis timing threshold in the cell membrane, resulting in a shorter lysis time. On the other hand, a lower rate of holin expression will result in a longer lysis time. All else equal, the lysis-triggering threshold in all lysogens should not change. Western blot assays using membrane extracts of holin from all lysogens confirmed similar holin levels in the membrane (data not shown). As a preliminary test, we estimated the lysis times for each strain by measuring OD_600_ over time of an induced culture of lysogens. Replacing the unusual SD sequence with the consensus sequence significantly lowered the lysis time compared to the S105 lysogen (CS, Fig. 1C). On the contrary, altering the SD sequence by a single nucleotide substitution, destabilizing or stabilizing the stem loop structures all caused significant delays in lysis times. However, lysis timing was not affected when the single nucleotide substitution in the SD sequence was combined with the mutation destabilizing the stem-loop (ASD+DSL, Fig. 1C). These mutations may be compensatory to each other and therefore reverse the delay in lysis caused by each individually.

For single-cell studies, we used an experimental set up where cells can be recorded using a camera attached to a microscope with a heated stage and a temperature-controlled perfusion chamber (Fig. 2A) (8). Fresh media flowed through the perfusion chamber to allow growth of immobilized cells in a continuous culture system. Cells grow normally under these conditions, and a temperature spike was used to induce the lytic cycle. Cell growth was visualized as an increase in cell length, which ends in lysis following induction (Fig. 2A). The recorded lysis times of single cells were used to analyze lysis time distributions (Fig. 2B and 4A) and calculate the mean (Fig. 2C) and noise (Fig. 4B). Compared to the S105 lysogen, the mean lysis time was observed to be shorter only for the lysogen containing the consensus sequence, while it was longer in all other cases (Fig. 2C).

### Effects of Tetracycline on Holin Translation

Translation efficiencies may be altered in the presence of antibiotics that affect translation. Tetracycline is an antibiotic that binds to the 30S ribosomal subunit and inhibits translation elongation (20). Sub-lethal doses for tetracycline were tested for their effect on lysis timing (Fig. 3). Most sub-lethal doses of tetracycline significantly reduce bacterial growth rate except at a 0.06 μg/mL concentration (inset, Fig. 3). As expected, lysis timing increased with increased tetracycline concentration. Intriguingly, tetracycline at a concentration of 0.06 μg/mL showed a significantly faster lysis timing compared to lysis timing in the absence of tetracycline. Tetracycline passively diffuses through the outer membrane porins OmpF and OmpC and accumulates in the periplasm (23–25). Transport into the cytoplasm is partially energy dependent and involves passive diffusion, phosphate bond hydrolysis, and the proton motive force (26–28). These mechanisms may induce changes in membrane permeability and affect the lysis process. We also speculate that feedback regulation of ribosome synthesis partially compensates for the slight decrease in the levels of available ribosomes following tetracycline inhibition (29). Thus, translation inhibitors such as tetracycline elicit an upregulation of ribosomal gene expression to counteract the effects of such antibiotics (30–32). This explains the faster lysis due to upregulation of ribosomes in the presence of very low doses of tetracycline. However, at higher tetracycline doses, the upregulation is insufficient due to very low levels of available ribosomes.

**Figure 3.**
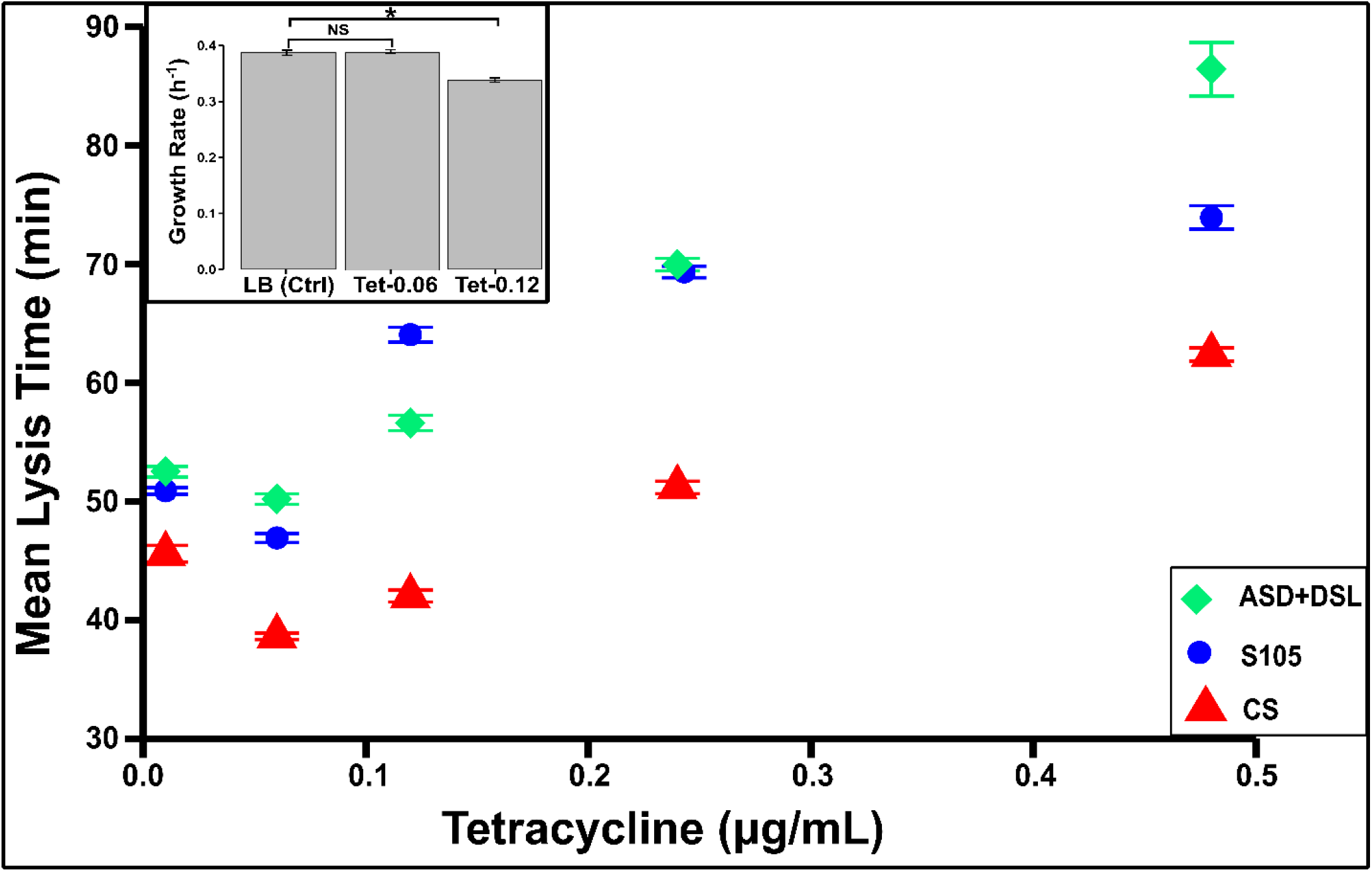
Effect of sub-lethal doses of tetracycline on mean lysis time. S105, CS, and ASD+DSL lysogens were tested for the effects of sublethal tetracycline doses on single-cell lysis timings. Each data point is the estimated mean lysis times of 200–300 cells. The mean drops initially and continues to rise as the tetracycline dosage increases. Error bars, 95% CIs after bootstrapping (1,000 replicates). Inset, bacterial growth rate comparisons in the presence of 0.06 and 0.12 µg/mL tetracycline. Error bars, mean ± S.E.M; NS, not significant. P values that are significant for a two-tailed test of less than 0.05 are denoted with an asterisk (n = 12).

### Modeling Lysis Timing in Individual Cells

To capture changes in the timing of lysis in response to translational perturbations, we borrowed stochastic frameworks previously used to model timing of intracellular events at the single-cell level that take into account the inherent noise in gene expression (33– 41). More specifically, after the onset of transcription from λ’s late promoter, intracellular holin concentration builds up over time as per a stochastic model of gene expression, and lysis occurs when holin levels reach a prescribed threshold level (1,2,42). The time taken from the onset of holin expression to holin levels reaching the lysis threshold was mathematically described as a first-passage time problem (42). We refer to this time as *FPT*—a random variable quantifying cell-to-cell difference in timing—and the lysis time is given by *FPT* plus 15 mins, with 15 mins being an approximate time delay between lysis induction and the start of holin expression (43,44). Previous work has derived the following formulas for the mean and variance in *FPTs* (denoted by ⟨*FPT*⟩ and 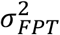, respectively) (9):

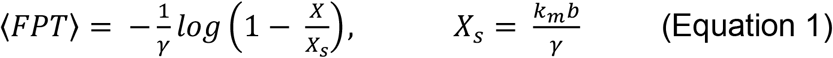

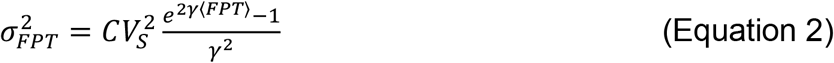

In these formulas, *k*_*m*_, *b* and *γ* describe the holin concentration buildup dynamics: *k*_*m*_ is the transcription rate from λ’s late promoter, *b* is the holin translational burst size and is directly related to the rate of holin translation from its mRNA, and *γ* is the rate of holin dilution. Parameter *X* is the threshold level of holin needed for host cell lysis to occur and parameter 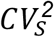 quantifies the steady-state noise in holin expression. We refer the reader to Kannoly et. al (9) for details on the stochastic model and assumptions employed in the derivation of the above formulas.

Using Equation 2, one can obtain an expression for *CV*_*FPT*_, the coefficient of variation of *FPT* as

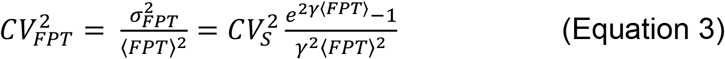

Interestingly, Equation 3 predicts the noise or the coefficient of variation of *FPT* to vary non-monotonically with the mean ⟨*FPT*⟩, and is minimized when *γ*⟨*FPT*⟩ ≈ *0*.*8* (9). For example, when *γ*⟨*FPT*⟩ ≪ 1, then 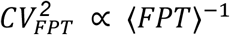, and *CV*_*FPT*_ decreases with increasing ⟨*FPT*⟩. However, for *γ*⟨*FPT*⟩ ≫ 1, *CV*_*FPT*_ increases with increasing ⟨*FPT*⟩ resulting in an overall U-shaped profile for the noise in timing with respect to the mean timing. This theoretical prediction was verified experimentally by altering the holin amino acid sequence to increase or decrease the lysis threshold *X* which yielded a six-fold change in the mean lysis timing (9).

While the derivation of the noise in *FPT* in Equation 3 is based on a small-noise approximation, this approximation has been relaxed in more recent work resulting in the derivation of exact *FPT* statistics for a class of stochastic gene expression model (45). Moreover, the first-passage time framework to model single-cell event timing has been extended in several directions, such as, the inclusion of extrinsic noise in gene expression (46), characterizing the impact of feedback regulation on timing (47,48), considering parameter fluctuations, for example, the threshold level itself varies across cells (49), and considering the role of decoys in regulating *FPT* (50). The latter extension is particularly important to understand the role of antiholin that sequesters and inactivates holin (16,51).

We first fitted Equation 3 to the data of noise in lysis timing across translation perturbations. These perturbations via site-directed mutagenesis or using non-lethal doses of tetracycline can both decrease or increase ⟨*FPT*⟩ via changes in the parameter *b* in Equation 1. After fixing the dilution rate *γ* = *1*.*035 hr*^−1^ (which corresponds to a 40 min *E. coli* cell-doubling time), 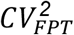 decreased with increasing ⟨*FPT*⟩ for the range of means lysis times considered in Fig. 4B. However, this decrease was quite shallow: only a 15-20% change in 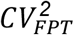 as ⟨*FPT*⟩ varied from 20 to 50 mins and was unable to explain the three-fold change in 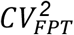 seen in the data (Fig. 4B). We next expanded Equation 3 to

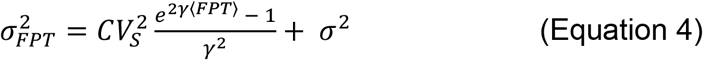

where the additional variance *σ*^*2*^ corresponds to either measurement noise or intercellular fluctuations in the time delay between lysis induction and the start of holin expression. Equation 4 yields the following formula for the coefficient of variation

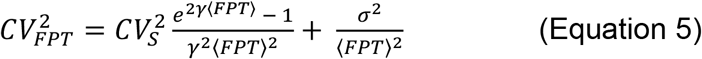

that is then fitted to the corresponding 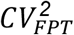 data as the mean ⟨*FPT*⟩ is varied through translational efficiency perturbations. Since the dilution rate *γ* = *1*.*035 hr*^−1^ is fixed, the fitting is done by varying the parameters *CV*_*s*_, *σ* in Equation 5 and performing a least-square optimization in Microsoft Excel using the Solver toolbox. The fitted Equation 5 is shown in Fig. 4B with parameter estimates *CV*_*S*_ = *2*.*6%* and *σ* = *0*.*06*.

**Figure 4.**
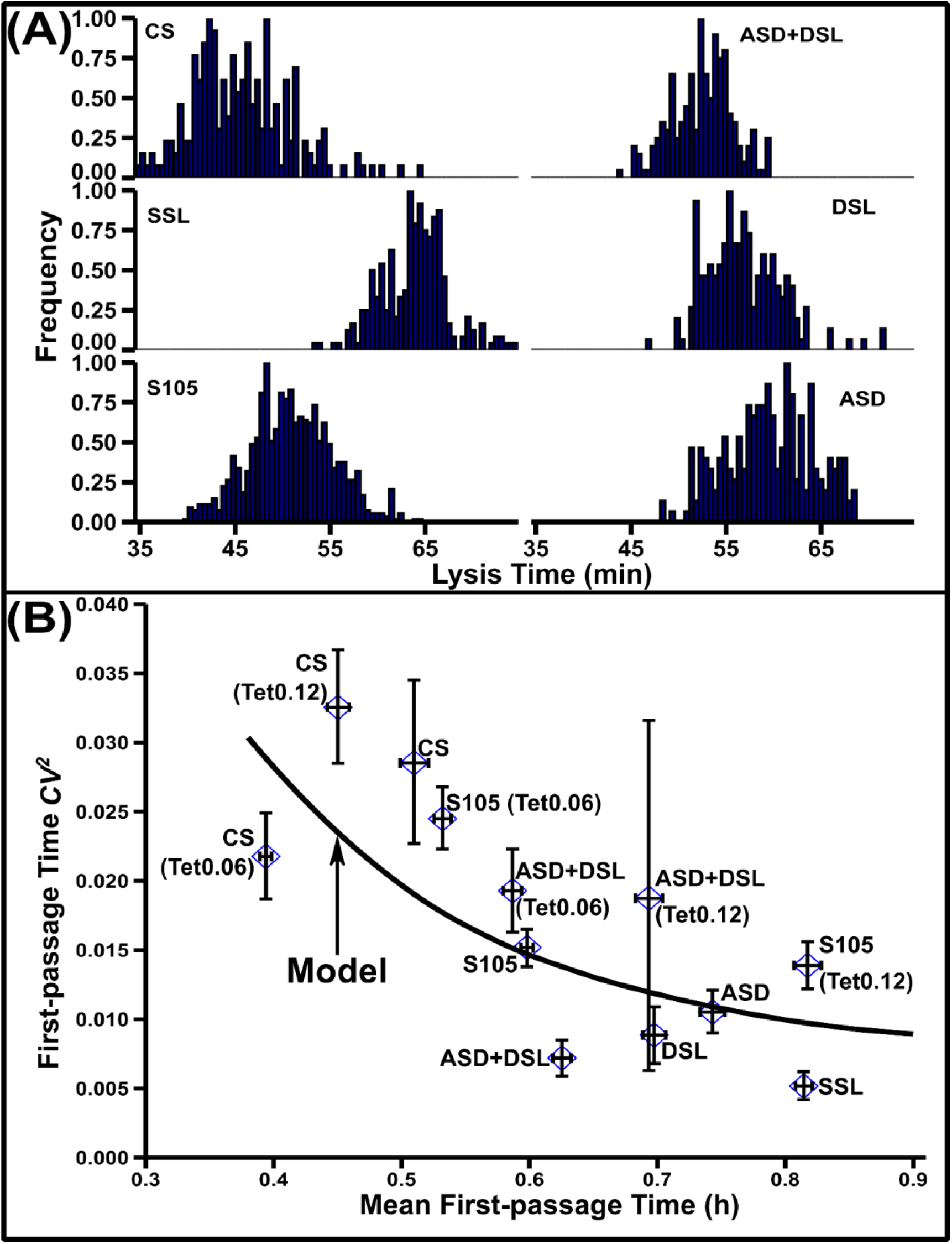
Effect of translation efficiency on lysis time noise. (A) Lysis time distributions for lysogens with different holin translational efficiencies. (B) The noise in first-passage time (FPT) is calculated using the coefficient of variation squared (*CV*^*2*^) and then plotted against mean FPT for S105, CS, ASD, DSL, ASD+DSL, and SSL lysogens. These lysogens show changes in FPT and the *CV*^*2*^ as predicted by the model (black line, Equation 5). Equation 5 was fitted to the data employing a 40-min doubling time for *E. coli* and fitting parameters *CV*_*S*_ = *2*.*6%, σ* = *0*.*06*.

## Conclusion

Previous studies concluded that holin structure and function has not only evolved to trigger lysis at an appropriate time but also to increase precision in lysis timing (5,9). It was further demonstrated that optimum lysis times that minimizes noise are correlated with improved phage fitness in a quasi-continuous culture (7). Our results here reveal that translational regulation of holin is another potential mechanism that λ may use to optimize lysis timing. Our results are supported by studies of *Bacillus subtilis*, which showed that increased translational efficiency is the predominant source of increased phenotypic noise (52). Similar outcomes were observed in theoretical studies of gene expression (53,54).

Replacing the unusual SD sequence in the holin gene with a consensus sequence improves translational efficiency causing faster lysis but also significantly increases noise (Fig. 4A and B). A shorter lysis time also reduces the burst size, thus affecting phage fitness. The presence of the unusual SD (UAAG) sequence instead of a sequence more similar to the typical G-rich sequences found in *E. coli* suggests a strong selection towards a weaker SD sequence (55). This observation is further supported by the delay in lysis observed in the ASD strain with an A-rich SD (Fig. 2C). However, the proteins translated downstream of holin, namely, endolysin (GCCGG) and spanin (GAGAG), appear to have stronger G-rich SD sequences indicating higher rates of expression compared to holin. Additionally, mutations in the stem loop structure always result in longer lysis timings. Thus, depending on the ecological challenges, lysis timing can be optimized by acquiring mutations in the translation regulatory regions. In summary, we demonstrated that holin translation efficiency and threshold independently impact mean lysis times and noise.

